# Explicit feedback and instruction do not change shoulder muscle activity reduction after shoulder fixation

**DOI:** 10.1101/2020.03.25.008466

**Authors:** Rodrigo S. Maeda, Julia M. Zdybal, Paul L. Gribble, J. Andrew Pruszynski

**Affiliations:** Brain and Mind Institute, Western University, London, Ontario, Canada; Robarts Research Institute, Western University, London, Ontario, Canada; Dept. of Psychology, Western University, London, Ontario, Canada; Dept. of Physiology and Pharmacology, Western University, London, Ontario, Canada

**Keywords:** Motor learning, arm dynamics, voluntary movements, visual feedback, explicit strategies

## Abstract

Generating pure elbow rotation requires contracting muscles at both the shoulder and elbow joints to counter torques that arise at the shoulder when the forearm rotates (i.e., intersegmental dynamics). Previous work has shown that human participants learn to reduce their shoulder muscle activity if the same elbow movement is performed after the shoulder joint is mechanically locked, which is appropriate because locking the shoulder joint eliminates the torques that arise at the shoulder when the forearm rotates. However, this learning is slow (i.e., it unfolds over hundreds of trials) and incomplete (i.e., shoulder activity is not fully eliminated). Here we investigated whether and how the addition of explicit strategies and biofeedback modulate this type of learning. Three groups of human participants (N = 55) performed voluntary pure elbow rotations using a robotic exoskeleton that permits shoulder and elbow rotation in a horizontal plane. Participants did the task with the shoulder free to move (baseline), then with the shoulder joint locked by the robotic manipulandum (adaptation), and then with the shoulder free to move again (post-adaptation). The first group of participants performed this protocol and received no instructions about what to do after their shoulder was locked. The second group of participants received visual feedback about their shoulder muscle activity after each movement and was instructed to reduce their shoulder activity to zero. The third group of participants also received visual biofeedback, but it was removed part way through the experiment. We found that, although all groups learned, the rate and magnitude of learning was not reliably different across the groups. Taken together, our results suggest that learning new arm dynamics, unlike other motor learning paradigms, unfolds independent of explicit instructions, biofeedback and task instructions.

## Introduction

Motor learning is complicated by the presence or absence of explicit cognitive strategies. Although previous work has demonstrated that explicit instructions can influence the ability of human participants to learn new movement tasks (Mazzoni and Krakauer 2006; Taylor et al. 2014), participants are also thought to engage in different strategies even when no such instructions are given (McDougle and Taylor 2019). Such variability in strategies might contribute to the often large variability observed in the rate and amplitude of learning across participants and tasks (Mazzoni and Krakauer 2006; Smith et al. 2006; Stark-Inbar et al. 2017). The effects of explicit instructions, however, have only been described in studies in which motor learning unfolds as a function of movement errors generated in the task, and in learning that takes place within a short number of trials.

We have recently showed that people learn to reduce shoulder muscle activity when generating pure elbow movements following mechanical fixation of their shoulder joint (Maeda et al. 2018). In this context, learning to reduce shoulder muscle activity is efficient because fixing the shoulder joint eliminates the torques that arise at the shoulder because of forearm rotation. Such learning takes place even when participants are not informed about the manipulation and without substantial kinematic errors in the task, which are two key pieces of information known to drive motor learning (Krakauer et al. 2019). Importantly, such learning unfolds very slowly, on a timescale of many hundreds of trials, and is incomplete even in that timeframe. Here we investigated whether and how the addition of explicit strategies and feedback modulate this type of learning. If this type of learning engages similar mechanisms known to be involved in traditional error-based motor learning paradigms, then we expect an influence of explicit instruction and feedback on the rate and amplitude of learning to reduce shoulder activity. An alternative possibility is that this type of learning is mediated entirely by an implicit mechanism that is independent of high-level instruction, and thus explicit components would not influence learning in this paradigm.

As in our previous work (Maeda et al. 2018), three groups of participants performed elbow flexion and extension movements using a robotic apparatus. First, they did so with the shoulder joint of the robot free to rotate (baseline phase). We then locked the shoulder joint of the robotic manipulandum, and participants repeated the same elbow movements (altered arm dynamics, adaptation phase). Lastly, we released the shoulder joint again and participants repeated the same elbow movements (post-adaptation phase). Using this protocol, we tested whether the rates and magnitude of learning these altered arm dynamics during reaching movements can be modulated by explicit strategies and feedback. One group of participants performed this protocol but received no instructions about what to do after their shoulder was locked, a replication of our previous studies (Maeda et al. 2018, 2020b). A second group of participants performed this same protocol with visual feedback about their shoulder muscle activity and was instructed to reduce their shoulder activity to zero. We found a reduction in shoulder muscle activity in both groups, but we found no differences in the rate and magnitude of learning across groups with the addition of strategies and feedback. A third group of participants performed the same protocol as the second group, but the explicit visual feedback was removed part way through the experiment. We found no reliable differences in shoulder muscle activity after visual biofeedback was removed, providing further evidence that visual feedback was not actively contributing to the reduction in shoulder muscle activity. Taken together, these results indicate that explicit strategies and feedback do not modulate learning of novel arm’s dynamics following shoulder fixation.

## Materials and Methods

### Subjects

Fifty-five healthy volunteers (aged 18-47, 30 female), participated in this study, each taking part in only one of the experiments described below. Participants self-reported that they were right-handed and that they had no history of neurological or musculoskeletal disorders. All participants signed a written letter of consent with the procedures approved by The Office of Research Ethics at The University of Western Ontario in accordance with the Declaration of Helsinki.

### Apparatus

Experiments were performed using a robotic exoskeleton (KINARM, Kingston, ON) that permits flexion and extension movements of the elbow and shoulder joints in the horizontal plane. Targets that participants had to move to, as well as their calibrated hand-aligned cursor, were projected from an LCD monitor onto a semi-silvered mirror in the same plane of motion. Direct vision of the participant’s arm was obstructed using a physical shield. The upper arm and forearm segments of the exoskeleton robot were adjusted to properly fit each participant’s arm and then filled with firm foam to ensure a tight coupling.

### Experimental Protocols

In all experiments, we used a similar protocol as described in Maeda et al. (2018). Fifty-five participants performed 20-degree flexion and extension trials with their shoulder joint free to move or fixed by the robotic manipulandum.

At the beginning of each trial, participants moved their hand to a home target, a red circle with 0.6 cm diameter. The home target position corresponded to when the participant’s shoulder and elbow joints were positioned at 10° and 60° (external angles), respectively (Figure 1, Task Setup). After maintaining their hand at this home target for a random period (250-500 ms, uniform distribution), a white goal circle with 3 cm diameter, was presented in a location that could be reached with a 20° pure elbow flexion movement. If the participant remained in this position for another random period (250-500 ms, uniform distribution), the hand feedback was extinguished, the goal target turned red, and the participant was instructed to start their movement. If participants failed to hold in these hold periods or failed to perform the movement within a 1 s time period, the hand feedback turned back on and the trial restarted. Participants were instructed to make their movements within a specific time frame (from start to end) as such the goal target turned green when movement time (measured as time from exiting the home target to entering the goal target) was between 100 and 180 ms, orange if it was faster than 80 ms, otherwise it remained red indicating that a slow movement took place. No restrictions were placed on movement trajectories. In addition to timing constraints between targets, participants were instructed to remain at the goal target for an additional 500 ms to finish a trial. After a random period between 0-1 s with uniform distribution, the goal target became a new home target with 0.6 cm diameter, and the same procedure was repeated but for an extension movement.

**Figure 1:**
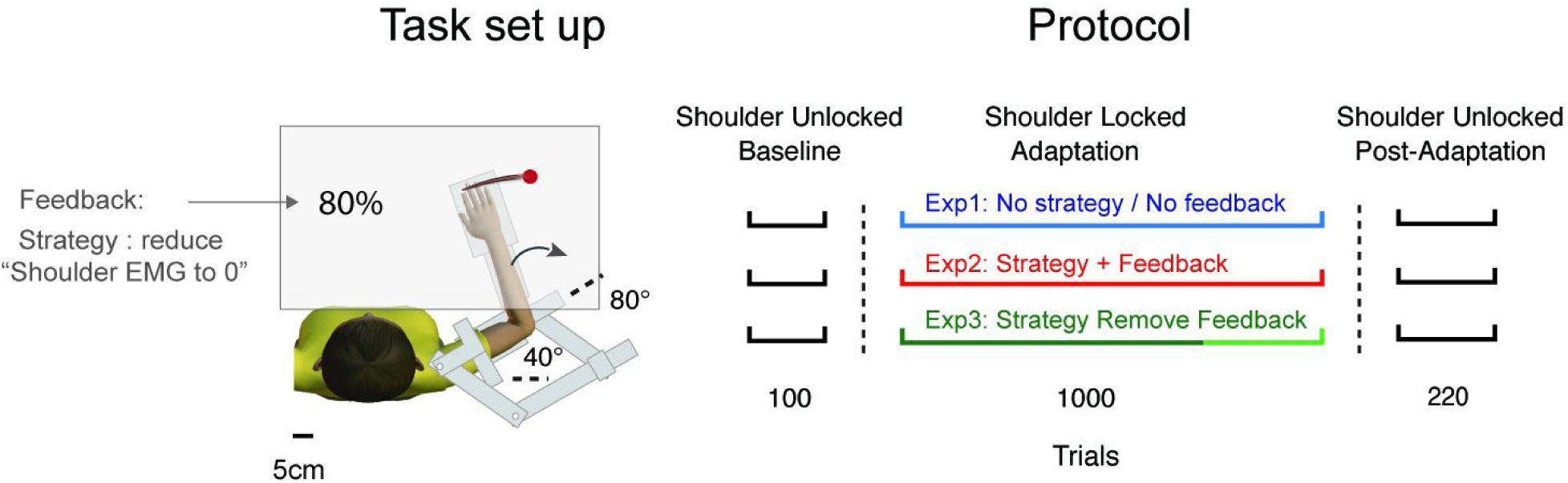
Experimental setup and protocol. For all experiments, participants used a robotic exoskeleton to perform twenty-degree flexion and extension trials with their shoulder joint of the robot free to move (black hand traces — left panel) or fixed (red hand traces— left panel). In Experiment 2 and 3, participants were given visual percentage feedback about their shoulder muscles and were instructed to reduce their shoulder activity to 0% (left panel). All participants first performed 100 elbow flexion and extension reaches with the shoulder joint free to move (baseline). We then mechanically locked the shoulder joint by the robotic manipulandum, and participants performed 1000 elbow flexion and extension movements (adaptation). Lastly, we released the shoulder joint and participants performed 220 elbow flexion and extension reaches with their shoulder joint again free to rotate (post-adaptation) (right panel). Instructions and availability of online visual biofeedback of the shoulder muscles varied between experiments. In the adaptation block of Experiment 1, participants received no strategy or biofeedback. In the adaptation block of Experiment 2, participants were given a visual percentage feedback about their shoulder muscle activity on the screen and they were instructed to reduce their percentages to zero. In the adaptation block of Experiment 3, participants were given the same instructions and visual percentage biofeedback, but this biofeedback was removed after 700 trials (right panel).

All experiments lasted about 2 hours. Participants were given breaks to rest when requested. Prior to the experiment, participants were given time to practice the task until they comfortably achieved an ∼80% success rate.

### Experiment 1: No Strategy or Feedback

Fifteen participants performed three blocks of trials for this experiment (Figure 1, Protocol). In the first block of trials, participants performed 100 elbow flexion and extension movements with their shoulder free to move (baseline). In a second block of trials, we mechanically locked the shoulder joint by the robotic manipulandum, and participants performed 1000 elbow flexion and extension reaches (adaptation). In this block of trials, participants were informed that their shoulders would be locked but they were told that this would not affect their ability to perform the task. In the third block of trials, participants performed 220 elbow flexion and extension reaches with their shoulder joint again free to move (post-adaptation).

### Experiment 2: Strategy and Feedback

Fifteen participants performed the same three blocks of trials for this experiment as in Experiment 1 (Figure 1, Protocol). However; in the adaptation block, participants were given a visual percentage feedback (see below) about their shoulder muscle activity on the screen and they were instructed to reduce this value to zero. They were told that this could reduce this value by reducing their shoulder muscle activation when performing the movement (Figure 1, Task set up).

Ten additional participants also performed a version of this experiment where they received the same instructions except that their shoulder joint was never locked (control experiment) (Figure 1, Protocol). This served as a control to quantify how much reduction in shoulder muscle activity reflects the explicit instruction and biofeedback as well as repeated movements rather than the shoulder locking manipulation per se.

### Experiment 3: Strategy and Removed Feedback

Fifteen participants performed the same protocol with the same instructions as in Experiment 2 (Figure 1, Protocol). However, after 700 trials in the adaptation block the visual percentage feedback was removed from the screen and participants performed the last 300 reaches without receiving any feedback.

### Visual Percentage Feedback

In Experiments 2 and 3, participants were given visual percentage feedback of their shoulder muscle activity on the screen in front of them (Figure 1, Task set up). This percentage was representative of the amount of muscle activity recorded from the posterior deltoid (PD) muscle during an elbow extension movement and was computed relative to baseline trials. Specifically, participants first completed the first baseline block of trials, the posterior deltoid data was extracted, rectified and normalized by baseline levels for each elbow extension trial. This baseline data was then averaged during the interval of -100ms to +100ms centered on when the hand left the home target (movement onset). This average was considered as 100% in the subsequent adaptation block. In the adaptation phase of these experiments, the average muscle activity from the PD muscle (−100ms to +100ms centered on when the hand leaves the home target) of each participant after each extension trial was compared to their own average muscle activity from the baseline trials and presented on the screen.

### Kinematic recordings, EMG recording and Analysis

Movement kinematics (i.e., hand position and joint angles) were sampled by the robotic apparatus at 1,000 Hz and then low-pass filtered (12 Hz, 2-pass, 4th-order Butterworth). In all experiments, data were aligned on movement onset, defined as 5% of peak angular velocity of the elbow joint (See Gribble and Ostry 1999; Maeda et al. 2017, 2018, 2020b).

Electromyographic signals from upper limb muscles were recorded with surface electrodes (Bagnoli-8 system with DE-2.1 sensors, Delsys, Boston, MA). These surface electrodes were coated with conductive gel and were placed on the surface of the skin aligned with the orientation of muscle fibers of five muscles: pectoralis major clavicular head (PEC), shoulder flexor; posterior deltoid (PD), shoulder extensor; biceps brachii long head (BB), shoulder and elbow flexor; brachioradialis (BR), elbow flexor; triceps brachii lateral head (TR), elbow extensor. The participants’ skin was abraded with rubbing alcohol prior to electrode placement. A ground electrode was placed on the participant’s left clavicle. EMG signals were amplified (gain = 10^3^), and then digitally sampled at 1,000 Hz. EMG data were then band-pass filtered (20–500 Hz, 2-pass, 2nd-order Butterworth) and full-wave rectified.

To compare the changes in amplitude of muscle activity over time and across different phases of the protocols, we calculated the mean amplitude of phasic muscle activity across a fixed time-window, -100 ms to +100 ms, relative to movement onset. These windows were chosen to capture the agonist burst of EMG activity in each of the experiments (See Debicki and Gribble 2005; Maeda et al. 2017, 2018, 2020a, 2020b).

Normalization trials prior to each experiment were performed and used to normalize muscle activity such that a value of 1 represents a given muscle’s mean activity when countering a constant 1 Nm torque (See Maeda et al. 2017, 2018; Pruszynski et al. 2008; Weiler et al. 2016). Data processing was performed using MATLAB (r2018a, Mathworks, Natick, MA). For simplicity, here we only report the learning profiles of extension movements; however, the results are similar for flexion (See Maeda et al. 2018).

### Statistical Analysis

We used repeated measures one-way ANOVA to compare each subject’s shoulder muscle activity at baseline to shoulder muscle activity after adaptation and again in the post-adaptation phase. We then used a one-way ANOVA to compare the amplitudes of learning after the adaptation phases across each experiment. We also used a one-way ANOVA to compare the initial rates of learning in each experiment by computing the mean shoulder activity across the first 20 bins of trials in the adaptation phase (Krakauer et al. 2005). This measure indicates how fast the subjects decreased shoulder activity. Where applicable we used Tukey post-hoc tests. All statistical analyses were performed in R v3.2.1. We considered experimental results to be statistically significant if the corrected p-value was < 0.05.

## Results

Participants (N=55) quickly met the imposed accuracy and speed of the task such that they were able to achieve >90% success rates within 5 min. of practice. Although no restrictions were placed on movement trajectories, participants did generate their movements with an arc-like trajectory, as a result of moving predominantly the elbow joint Figure 2 A,C,E (Maeda et al. 2017, 2018, 2020a, 2020b). We found substantial shoulder agonist muscle activation prior to movement onset in all experiments as required to compensate for the torques that arise at the shoulder when the forearm rotates about the elbow joint Figure 2 B,D,F (Gribble and Ostry 1999; Maeda et al. 2017, 2018, 2020a, 2020b). We then mechanically locked the shoulder joint of the robotic manipulandum, which cancelled the torques that arise at the shoulder with forearm rotation and removes the need to activate the shoulder muscles. Locking the shoulder had no effect on participants’ ability to perform the task and they continued to achieve >90% success rate.

**Figure 2:**
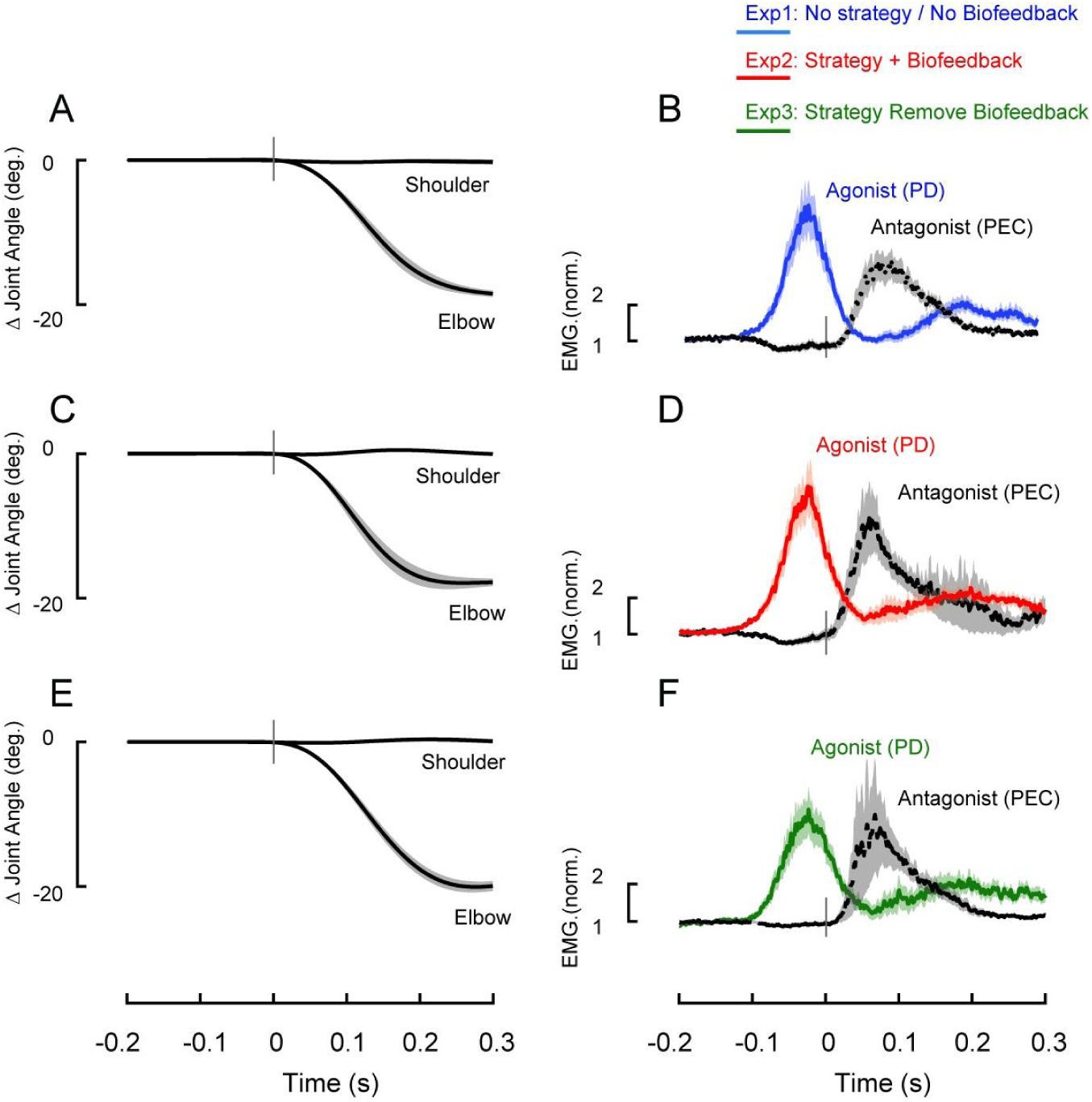
Compensating for intersegmental dynamics during voluntary elbow reaches. A. Average kinematics of the shoulder and elbow joints during elbow extension movements in the baseline trials of Experiment 1 (shoulder joint free to move). Shaded areas represent the standard error of the mean (SEM). Data are aligned on movement onset. B. Average agonist (PD) and antagonist (PEC) muscle activity associated with movement in A. Shaded areas represent SEM. Data are aligned on movement onset. EMG data normalized as described in Methods. C-D, Same format as A-B but from Experiment 2. E-F, Same format as A-B but from Data associated with the Experiment 3.

### All Experiments: Reducing shoulder muscle activations with shoulder fixation

First, we confirmed that the nervous system learns to reduce shoulder muscle activity when the shoulder is mechanically fixed (Maeda et al. 2018). Figure 3A-B shows the mean shoulder extensor (PD) muscle activity in a fixed time window (−100 to 100ms centered on movement onset, see Methods) over elbow extension bins of trials in all experiments before locking the shoulder joint (i.e., baseline phase), with the shoulder locked (i.e., adaptation phase) and after unlocking the shoulder joint again (i.e., post-adaptation phase). To compare shoulder agonist muscle activity across these learning phases (last 25 trials in the baseline trials, adaptation and post-adaptations phases), we performed one-way ANOVAs for each experiment. We found a reliable effect of phase in each experiment (Experiment 1: F_2,28_ = 17.3, p < 0.0001, Figure 3A and 3C; Experiment 2: F_2,24_ = 16.5, p < 0.0001, Figure 3A and 3C; Experiment 3: F_2,24_ = 9.3, p < 0.0009, Figure 3B and 3C). Tukey post-hoc tests showed a reliable reduction of shoulder muscle activity with shoulder fixation from baseline trials in each experiment (Experiment 1: 31%, p < 0.0001; Experiment 2: 35%; p < 0.0001; Experiment 3: 24%; p < 0.0001), which returned to baseline levels after releasing the shoulder joint again (Experiment 1: p = 0.334; Experiment 2: p = 0.1; Experiment 3: p = 0.32). We found no reliable effect of phase in a control experiment (see Methods) where participants never experienced shoulder fixation but performed the same task with the same instruction and feedback as in Experiment 2 (F_2,18_ = 2.89, p = 0.08, Figure 3E and 3F). Importantly, shoulder muscle activity at the end of the adaptation phase in Experiment 2 was reliably smaller than at the equivalent point in the control experiment (t = -2.69, df = 20.9, p = 0.01, Figure 3G), indicating that explicit instructions, biofeedback and extensive practice were not the sole drivers of shoulder muscle activity reductions in Experiments where the shoulder was mechanically fixed.

**Figure 3:**
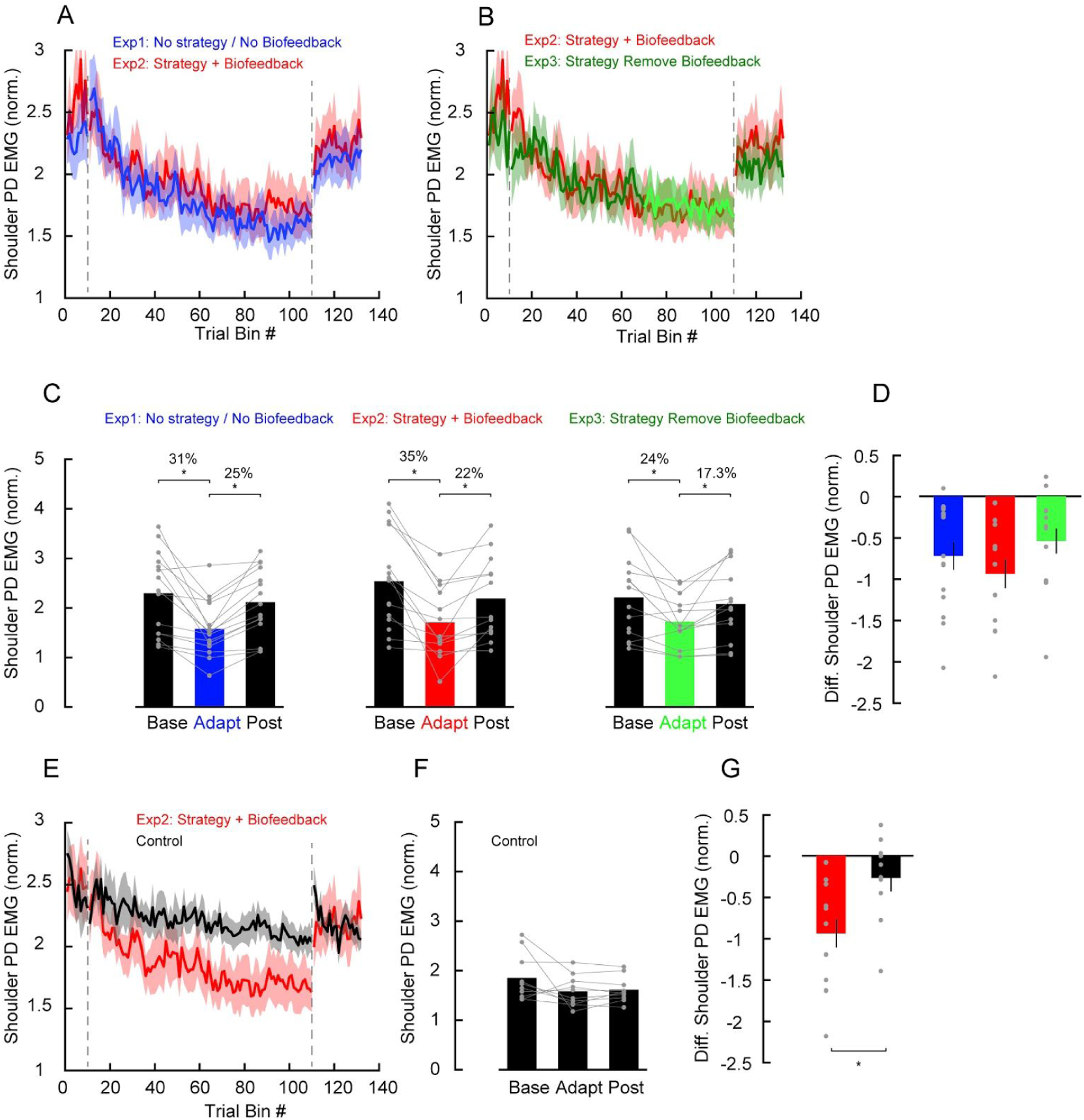
Effects of instruction and feedback on the adaptation of shoulder muscle activity following shoulder fixation. A-B. Average PD muscle activity in a fixed time window (−100 to 100 ms relative to movement onset of elbow extension trials) before locking the shoulder joint (i.e., baseline phase), with the shoulder locked (i.e., adaptation phase) and after unlocking the shoulder joint again (i.e., post-adaptation phase). Each phase is separated by vertical dashed lines. Each data bin is the average of 5 trials. Shaded areas represent SEM. EMG normalized as described in Methods. Blue traces represent data from Experiment 1, with no instruction or biofeedback. Red traces represent data from Experiment 2, with instruction and biofeedback. Green traces represent data from Experiment 3, with instruction and biofeedback but where biofeedback was removed near the end of the adaptation phase (light green). C. Average of the last 25 trials in each phase of the protocol for each Experiment. Each grey dot represents data from a single participant. Asterisks indicate reliable effects (p<0.05, see main text). D. Average of the difference of shoulder PD muscle activity (adapt-baseline) of the data associated with panel C. Error bars represent (SEM). Each grey dot represents data from a single participant. E-F data of the control Experiment and Experiment 2, with instruction and biofeedback, in the same format as in A and C. G. Average of the difference of shoulder PD muscle activity (adapt-baseline) of the data associated with control Experiment and Experiment 2, with instruction and biofeedback in panel E. Error bars represent (SEM). Each grey dot represents data from a single participant.

We found no corresponding reduction in elbow extensor (TR) muscles over trials with shoulder fixation (Experiment 1: F_2,28_ = 5.7, p = 0.008—with Tukey post-hoc test showing no reduction from baseline levels (p = 0.62); Experiment 2: F_2,24_ = 0.18, p = 0.83; Experiment 3: F_2,24_ = 0.67, p < 0.5).

### Comparing amplitudes and rates of shoulder activity reduction

To test whether the amplitude of shoulder muscle activity was different between experiments after learning with shoulder fixation, we performed a one-way ANOVA on the mean normalized shoulder muscle activity of the last 25 trials in the adaptation phase (middle bars in Figure 3 C). We found no reliable differences between Experiments 1 (with no instruction and feedback), 2 (with instructions and feedback), and 3 (with instructions and feedback in the beginning but with feedback removal in the middle of the adaptation phase) (F_2,41_ = 0.25, p = 0.78, η2 = 0.01). Similarly, we found no reliable differences in the amplitude of the reduction of shoulder extensor activity (adapt-baseline) across the three experiments (one-way ANOVA, F_2,41_ = 1.27, p = 0.29, η2 = 0.06, Figure 3 D). Lastly, to compare the initial rate of shoulder activity reduction after shoulder fixation, we computed the mean shoulder muscle activity for the first 100 trials (20 bins of trials) in the adaptation phase for each experiment (Krakauer et al. 2005). A one-way ANOVA revealed no reliable difference in learning rates early in the adaptation phase across the three experiments (F_2,41_ = 0.29, p = 0.75, η2 = 0.013, Figure 4 A-B).

**Figure 4:**
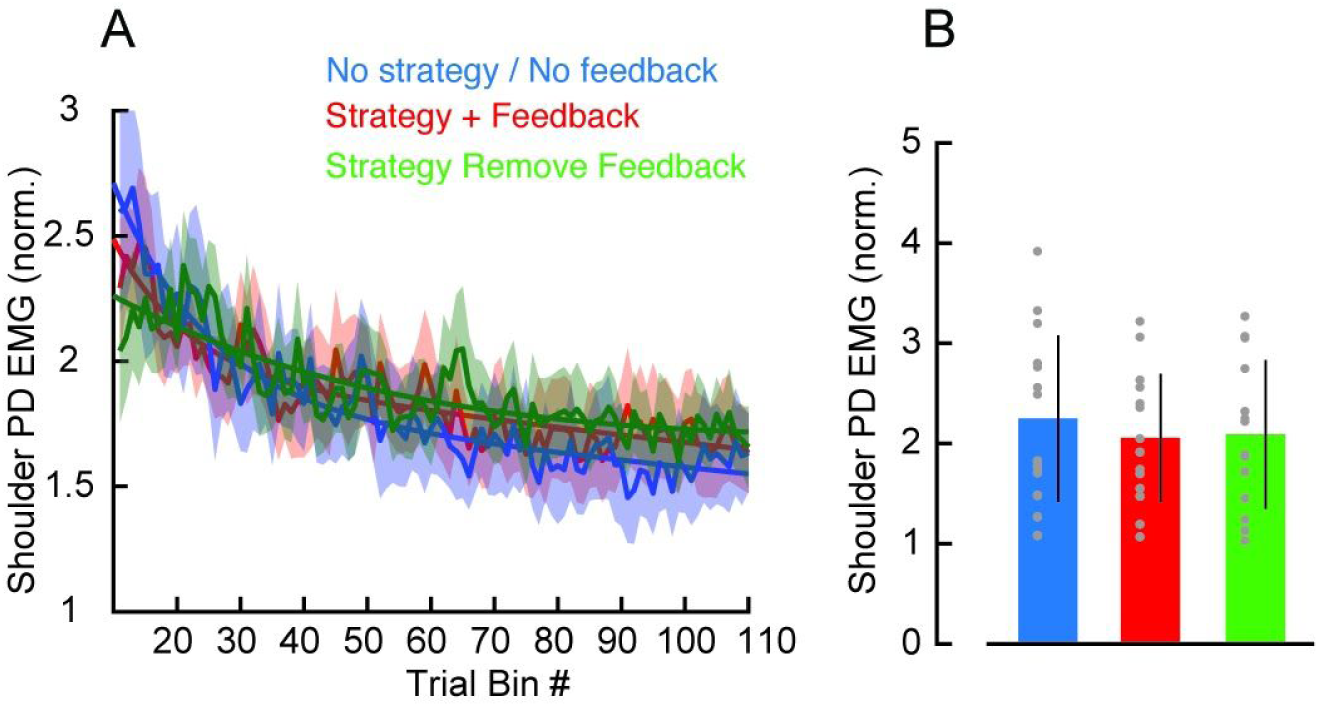
Rates of shoulder muscle activity reduction following shoulder fixation with different instructions and feedback. A. Average PD muscle activity in a fixed time window (−100 to 100 ms relative to movement onset) during elbow extension trials with the shoulder locked (i.e., adaptation phase). Each data bin is 5 trials. Shaded areas represent SEM. Blue traces represent data from the experiment with no instruction or biofeedback. Red traces represent data from the experiment with instruction and biofeedback. Green traces represent data from experiment with instruction and biofeedback but that the biofeedback was removed in the adaptation phase. Lines represent the result of an exponential fit to the averaged profiles. B. Average PD muscle activity in a fixed time window (−100 to 100 ms relative to movement onset) in trials early in the adaptation phase (first 20 bins) associated with the experiments in panel A. Error bars represent (SEM). Each grey dot represents data from a single participant.

## Discussion

In Experiments 1 and 2, we tested whether learning to reduce shoulder muscle activity for pure elbow rotations after shoulder fixation is influenced by the presence or absence of explicit instruction and biofeedback about the task. We found no differences in the rate and magnitude of the reduction in shoulder muscle activity after shoulder fixation despite adding strategies and biofeedback. In Experiment 3, we investigated whether the presence of explicit biofeedback was contributing to the ongoing reduction in shoulder muscle activity by removing it midway through the adaptation epoch. We found no significant differences in the reduction of shoulder muscle activity after feedback removal. Taken together, these results suggest that learning new arm dynamics following shoulder fixation unfolds independently of explicit task instructions and feedback.

### Reducing shoulder muscle activity after shoulder fixation with instruction and feedback

Many previous studies have investigated the role of explicit cognitive strategies in motor learning (Bond and Taylor 2015; de Brouwer et al. 2018; Christou et al. 2016; Mazzoni and Krakauer 2006; McDougle et al. 2015; Miyamoto et al. 2020; Poh et al. 2016; Rand and Rentsch 2015; Stark-Inbar et al. 2017; Taylor et al. 2014). Regardless of the type of learning studied (i.e., visuomotor rotation, forcefield, etc.), explicit strategies in these tasks quickly improved learning speeds (for review, see Krakauer et al. 2019). We previously reported that people learn to reduce shoulder activity after shoulder fixation, which unlike many other motor learning paradigms, unfolded slowly over hundreds of trials and it was still incomplete by the end of the testing session (Maeda et al. 2018, 2020a, 2020b). In addition, this learning took place without people knowing about this manipulation and in the absence of kinematic errors, as locking the shoulder joint restricts kinematics into an arc about the elbow joint. Here we tested whether and how much such learning could be modulated by giving participants explicit instructions and feedback to voluntarily reduce shoulder activity with shoulder fixation. We found no differences in learning with and without these explicit components, indicating that the presence of an explicit strategy does not lead to universal changes across motor learning tasks.

We provided participants with visual feedback of the amplitude of their muscle activity, as a percentage from baseline reaches, on a trial by trial basis with the shoulder locked and instructed participants to reduce those numbers to zero. We expected participants to use this direction and magnitude information, as an error signal in this task. Why was there no effect of visual feedback and task instruction in this type of learning? In error-based learning paradigms, instructions and error feedback are normally linked to the goal of the task. For instance, if a force field diverges reaching to the left of the goal target, people are instructed to aim or to perform the next reach to the right of the workspace in order to bring the hand in the target (Mazzoni and Krakauer 2006; Taylor et al. 2014). In our task, however, the instruction and feedback had no direct influence on the movement outcome but only on how efficiently the movement was generated. The main reason for this approach, however, was because this type of learning does not unfold as a function of errors. Rather, learning unfolds as a means to increase efficiency as shoulder muscle activity is not needed since mechanically fixing the shoulder joint cancels the interaction torques that arise at the shoulder when the forearm rotates.

Moreover, another possibility is that this learning is mediated by or dominated by an implicit mechanism (Mazzoni and Krakauer 2006; Miyamoto et al. 2020). Consistent with this notion, learning in our task tends to be slow and incomplete (Maeda et al. 2018, 2020b) and previous work has even supported the existence of a fixed synergy between shoulder and elbow muscles (Debicki and Gribble 2004; de Rugy et al. 2012). One important venue for future research is to test whether this is the case only for the timescale studied in these experiments or whether people would be able to use such instructions following extended periods of practice over many sessions.

### Neural structures for learning arm dynamics

Learning involving explicit cognitive strategies has been associated with engagement of the prefrontal cortex (PFC) (Asaad et al. 1998; Christou et al. 2016). Error-based learning has long been associated with the cerebellar-dependant adaptation that occurs in many forms of motor learning (Krakauer et al. 2019; Schlerf et al. 2012). Here we found no modulation of shoulder activity reduction with shoulder fixation by explicit task instructions and visual feedback, which suggest a different mechanism that might be in play for this type of motor learning. One possibility is that this learning happens at low levels subcortical or spinal processes, independently of these high level instruction mediated areas (Asaad et al. 1998; Christou et al. 2016; Floyer-Lea and Matthews 2004, 2005; Huberdeau et al. 2015; Shadmehr and Holcomb 1997; Taylor and Ivry 2014). For instance, the spinal cord is known for hosting neural circuits that generate patterns of muscle activity (Duysens and Van de Crommert 1998; Weiler et al. 2019). In addition, the spinal cord’s involvement in learning motor behaviours has been demonstrated in studies involving spinal cord transection, in which subsequent training improves locomotor activity (Brownstone et al. 2015; Harkema et al. 2012; Leblond et al. 2003). Thus, the reduction in shoulder activity might be a reflection of changes at the spinal level. This prediction could be confirmed in future studies investigating whether short-latency spinal responses change as a function of this type learning.

Our results have direct implications for people with musculoskeletal impairments (i.e., long-term injuries), neurological (i.e., stroke) or people that undergo joint immobilization for extended periods of time (i.e., bone fractures). For instance, rehabilitation approaches focusing on constraint-induced movement therapies was demonstrated to be beneficial for stroke patients when training of the unaffected limb occurs with a constraint in the unaffected limb (Dromerick et al. 2000; Krakauer 2006; Mark and Taub 2004). In addition, rehabilitation approaches also took advantage of EMG biofeedback (Giggins et al. 2013). Thus, knowing the timescale of the learning rates in this scenario, as well as the roles of instruction and feedback in this process may prove critical for the development of more efficient rehabilitation strategies.

### Limitations

There are limitations of our study that should be considered. First, although we found no reliable differences in the degree or rate of shoulder muscle activity adaptation across experiments, it is hard to prove the absence of an effect due to many intrinsic experimental factors, including sample size and number of repetitions. Note, however, that we tried to keep these experimental factors as close as possible to our previous studies (Maeda et al. 2018, 2020a, 2020b), and in particular to a study in which we did observe reliable differences across experiments (Maeda et al. 2020b). Second, the metric used as feedback to the participant (muscle activity amplitude) is a relatively noisy measurement. Thus, the value shown on any given single trial has substantial uncertainty with respect to the actual outcome and feedback uncertainty is known to slow down motor learning (Wei and Körding 2010). However, in several previous studies EMG metrics have been used for biofeedback and in these studies participants are able to modulate EMG given instructions. This kind of EMG-based biofeedback has also been demonstrated to be useful for rehabilitation and prosthetic control (Dosen et al. 2015; Giggins et al. 2013; Kim 2017; Radhakrishnan et al. 2008; Thompson and Wolpaw 2014; Woodford and Price 2007). Lastly, although it is tempting to say that the learning we describe happens implicitly, the absence of an effect of explicit cues in this experiment does not mean that people are not engaged in different strategies (McDougle and Taylor 2019). In fact, people were aware that the shoulder joint was locked and when it was locked. However, the main goal of this study was not to examine the effects of the explicit and implicit systems in isolation but instead to investigate whether additional explicit instructions and feedback can modulate this type of learning with shoulder fixation. One possibility for future work is to use approaches known to isolate implicit learning mechanisms, such as by gradually introducing shoulder fixation and/or introducing the manipulation without participants knowing about it, a condition previously studied for error-based learning paradigms (Orban de Xivry et al. 2011, 2013).

